# A computational strategy for rapid on-site 16S metabarcoding with Oxford Nanopore sequencing

**DOI:** 10.1101/2020.08.25.267591

**Authors:** Stefano M. Marino

## Abstract

The investigation of microbial communities through nucleotide sequencing has become an essential asset in environmental science, not only for research oriented activities but also for on-site monitoring; one technology, in particular, holds great promises for its application directly in the field: the Oxford Nanopore Technologies (ONT) MinION sequencer is a portable and affordable device, that produces long reads, with a remarkable sequencing output (in terms of bases/hour). One of the most common approaches in microbiological investigations through sequencing is the analysis of the 16S rRNA gene, known as 16S metabarcoding. Only recently the application of MinION has extended to 16S metabarcoding; to date, a limitation is still represented by the available computational protocols: due to the intrinsic, unique features of the technology ONT long reads cannot be adequately analyzed with tools developed for previous technologies (e.g. for Illumina). In this work a computational pipeline, specifically tailored to the usage of ONT reads in 16S metabarcoding, is developed, tested and discussed. This study is particularly addressed to on site evaluations, for environmental investigations or monitoring, where running time, costs and overall efficient usage of resources are particularly important.

## INTRODUCTION

Metagenomics can be defined as the study of genetic material recovered from the environment, with direct sampling. Since the last forty years, the advent of modern molecular biology techniques allowed for increasingly specific genetic analyses of complex samples, expanding our knowledge of ecological systems and their structure. In environmental analysis, molecular typing (e.g. via PCR, with targeted or random amplification; or by enzymatic restriction) (Maiden et al, 1998; Van Belkum, 2002; Sabat et al, 2013; Luna, 2016; Foley at al, 2009) opened for a highly specific detection of target DNA regions (such as marker genes, that can serve to trace a trait) and for their characterizations at the population level. In general, molecular methods have significantly expanded our ability to describe environmental microbiology, beyond the limitation of addressing only those cultivable bacterial taxa that can be studied by classical microbiological methods (Fredricks and Relman, 1996). The rise of nucleotide sequencing, and next generation sequencing (NGS) in particular, marked another significant step forward: besides the most upfront capabilities offered (e.g. genome sequencing), it significantly expanded the possibility of molecular typing, including population-wise characterization of environmental samples. With NGS, the definition of metagenomics is often referred to the random ‘shotgun’ sequencing of microbial DNA, and successive computer-based classification, without selecting any particular gene. Instead, when the analysis is conducted on a particular gene (or target region), the term metabarcoding is often employed (Breitwieser et al, 2019). As for the latter, the 16S ribosomal RNA (16S) gene have been the most predominantly used molecular marker, for bacterial classification (Srinivasan et al, 2015). The bacterial 16S gene is ∼1500 bp, and contains conserved and variable regions, evolving at different rates (lower for conserved, higher for variable regions), meaning that the former regions are well suited for the design of universal primers (for amplification of 16S genes across different taxa); at the same time, variable regions are critical to reflect differences between species and thus provide the most useful information for taxonomic purposes (Clarridge, 2004). NGS for microbiological analyses can be applied to on-site characterization of samples, in environmental monitoring (Sabat et al, 2013; Pérez-Losada et al, 2018); this could apply to situations of emergency, from medical hazards (pathogens outbreak) to environmental stress (e.g. pollutants contamination, spills, etc); indeed, similar situations are increasingly becoming an urgent topic. In this respect, one of the currently available NGS technology clearly stands out: the Oxford Nanopore Technologies (ONT) MinION sequencer. This is a very portable (light, pocket size), affordable and energy efficient (limited power requirements to run) technology, while producing a significant sequencing output (Gigabases in hours). The MinION already found application in all the contexts discussed in previous paragraphs; however, up to very recently its usage in bacterial metabarcoding was negligible, with 16S metabarcoding relying mostly on Illumina sequencing technology. In respect to Illumina, MinION reads are considerably less accurate (around 90%, for 16 genes), but the technology allows the sequencing of the full-16S rRNA gene (while Illumina only targets some sub-regions, e.g. V3 and V4), a feature which increases the capacity for taxonomical classification; it has been shown that this compensates the higher error rate (Cuscó et al, 2018; Leggett et al, 2020). As previously highlighted, the MinION presents features that are ideal for fieldwork: besides the outstanding features of the sequencer *per se*, optimized protocols and portable equipment have been developed and tested for on-site analyses, including in extreme conditions and/or remote locations (Quick et al, 2016; Castro-Wallace et al, 2017; Arwyn et al, 2019; Maestri et al, 2019). Overall, the MinION holds great promises to become a reference tool for on-site environmental genomics. Currently, there is a need of dedicated computational tools for ONT sequencing applied to 16S metabarcoding, with most tools being developed for high accuracy reads (e.g. from Illumina technologies) (Santos et al, 2020). In this respect, Illumina benefits from a large community and several years of development. ONT MinION reads requires dedicated pipelines, not only to deal with the lower accuracy but also to handle the specific file formats (e.g. fast5) and to pipeline the different utilities (for basecall, barcoding, adapters removal, etc). Oxford Nanopore provides a specific tool for 16S metabarcoding, called Epi2me; this proprietary pipeline has been employed in some studies (Cuscó et al, 2018; Gonçalves et al. 2020), but presents important limitations (Santos et al, 2020), such as restricted usage (to ONT customers, and with proprietary software, e.g. a Desktop Agent software, requiring web access), no possibility to customize any of the search parameters (all fixed, configured by default) and providing an output for the taxonomic assignments that, while graphically intuitive, does not allow downstream computational analyses (e.g. reads assignments, diversity calculations, differential abundances of specific taxa, etc; basically, the output is an interactive, graphical representation of the taxonomy distribution of genera represented in the sample). Finally, while currently “for free” (albeit, restricted to ONT registered users), it should become “for fee” (based on a credit systems, see ONT website). For once, Epi2me could not have been used for an analysis like the one presented in this work. Here, a simple (in structure and usage) computational pipeline, termed 16S_ppm, for 16S metabarcoding with ONT sequencing is presented and tested. As mentioned before, the study and the computational approaches were designed specifically for an application in the context of on-site analyses, for a rapid response and with portable equipment (e.g. computationally, a standard computer that both powers the MinION, and runs the classification; e.g., while the sequencing is running).

## METHODS

### Source Material

Sequencing data for this study were taken from different sources, from recently published (e.g. as NCBI projects, or as peer reviewed publications) works. Two mocks were used, for validation purposes: the first is the mock HM-783D, a community of 20 different bacterial strains, characterized by 16S amplification and sequencing with ONT MinION (available through NCBI website, SRA code:SRR8029996). The second study refers to a mock community of 12 bacterial strains, called mock B12 (SRA:SRR8351023), relative to a metagenomics study with ONT MinION. Since the reads of this experiment were derived from “shotgun” metagenomics (and not with a specific 16S amplification), the reads related to 16S genes were “fished” among the full dataset (with Blast, as described below), thus creating a subset of the original data made only of reads with 16S (partial or complete) gene information (for a total of 1,828 16S reads). Finally, a third mock, also from a metagenomics study, was employed, called ZymoBIOMICS™ (Zymo Research Corp Instruction Manual ZymoBIOMICS; Gonçalves et al, 2020); this mock is composed of 20 bacterial species and was used for additional validation (see Supporting information). For the analyses of environmental samples, different data were retrieved from the NCBI SRA repository. A study based on 16S metabacarcoding with ONT of environment DNA was recently made available (NCBI website, project code:PRJNA554976; SRA: SRP214990), investigating different chemically stressed anaerobic digesters, sampled in Germany. Of these samples, one in particular (SRR9695976, with library code name “C3.1-20-8”) was used for various tests, discussed in the Results section. Additionally, a very recent (Gonçalves et al, 2020) study, on microbiota communities associated with salmon parasites from different locations, was considered for further evaluations (particularly, the effects of sub-sampling on the general results); the project code of this study is PRJNA598707 (see also Supporting information). Diversity analyses were done with two main metrics; the diversity in terms of species within one sample, was evaluated as alpha-diversity (with Shannon, As), with the following equation:

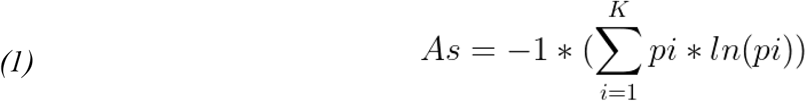

with *i* being the *i-th* taxon, and *K* is the number of taxa; the diversity in terms of species between two samples, was evaluated as beta-diversity Bray Curtis (Bc), with the following equation:

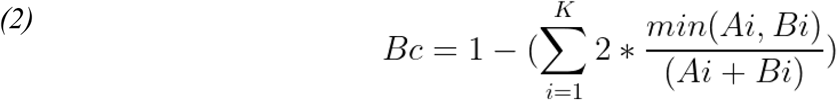

### Pipeline

*T*wo aspects were first considered: the choice of the reference database, and the choice of the aligner. First, the database of reference was as choice between GreenGenes (DeSantis et al, 2006), SILVA (Pruesse et al, 2007) (particularly useful if also other rRNA regions, e.g. 18S, are of interest), and NCBI 16 database (Federhen, 2015). Of them, NCBI 16S is the most curated and updated. GreenGenes is still widely used in 16S analyses with Illumina sequencing, as it is employed in the popular suite *qiime* (Caporaso et al, 2010); however, it is now becoming obsolete, as it has last been updated in 2013 (Balvociute and Huson, 2017). Conversely, the NCBI 16S database is more recent, actively maintained and regularly updated by NCBI (Balvociute and Huson, 2017), and it has become a reference in the Nanopore community (Santos et al, 2020), also because it was chosen by ONT as the reference database for their proprietary 16S analysis pipeline, called Epi2me. Last but not least, the NCBI 16S database is locally downloadable and can be easily edited and customized (which can be very useful, for some dedicated applications). For the aligner, different options were initially considered (Blast, LAST, minimap2); minimap2 (Li, 2018) is a mapper, a popular choice for mapping reads to reference genomes (with MinION, it is popular for metagenomics; Santos et al, 2020), but its usage as an aligner with a large reference set of 16S genes (rather than mapping the reads on a set of genomes; Kai et al, 2018) seems not adequate for the task (for once, the computational costs of mapping against all available bacterial genomes are significantly higher, than direct read-to-16S reference genes alignment; and would require a set up more in line with long and resource intensive servers, i.e. not the target of this study). As for the aligners: both Blast (Altschul et al, 1990) and LAST (Kielbasa et al, 2011) were initially tested, and ultimately the former was employed; in the tests, it provided a higher annotation rate, and better correlation with mock actual abundances (see Results section, and Supporting information, Fig. S2); moreover, testing LAST with customized parameters, as suggested in (Xia et al 2017), did not work well for this task: it did not improve the results, while implying much longer computations (up to 10 times slower, with mock data). As for the taxonomic assignment: ONT chose for its Epi2me pipeline to limit the analysis at the genus level; considering the accuracy at around 90% and the overall similarity of 16S genes (which can share an overall similarity > 95 % for different species), they consider the genus level as the safest option. The same view is shared in this work: while the 16S_ppm pipeline returns results at the species level (and, in some cases, with sufficiently accurate results), data interpretation has been mostly at the genus level. Blast versions employed included older (Blast 2.2.31) and recent (Blast 2.9.0) versions, to compare the effects on results, particularly the options “-max_target_seqs” and “-num_threads”. The Blast output was filtered for e-value (< 0.00001); for analyses of environmental samples (e.g. in Fig. 2) additional filtering were applied for coverage (> 60%) and % of sequence identity (> 70%). Then, for each read, after parsing all filtered alignments, the best hit (by Blast score) with the NCBI 16S database was extracted, and accordingly the taxonomic description was assigned. Before entering the pipeline for classification, adapters were removed from the fastq with Porechop (0.2.4), reads were filtered with Nanofilt (v 2.2.0), with options -q 7 and -l > 800 bp. A scheme of the pipeline is reported in Fig. 1.

**Figure 1.**
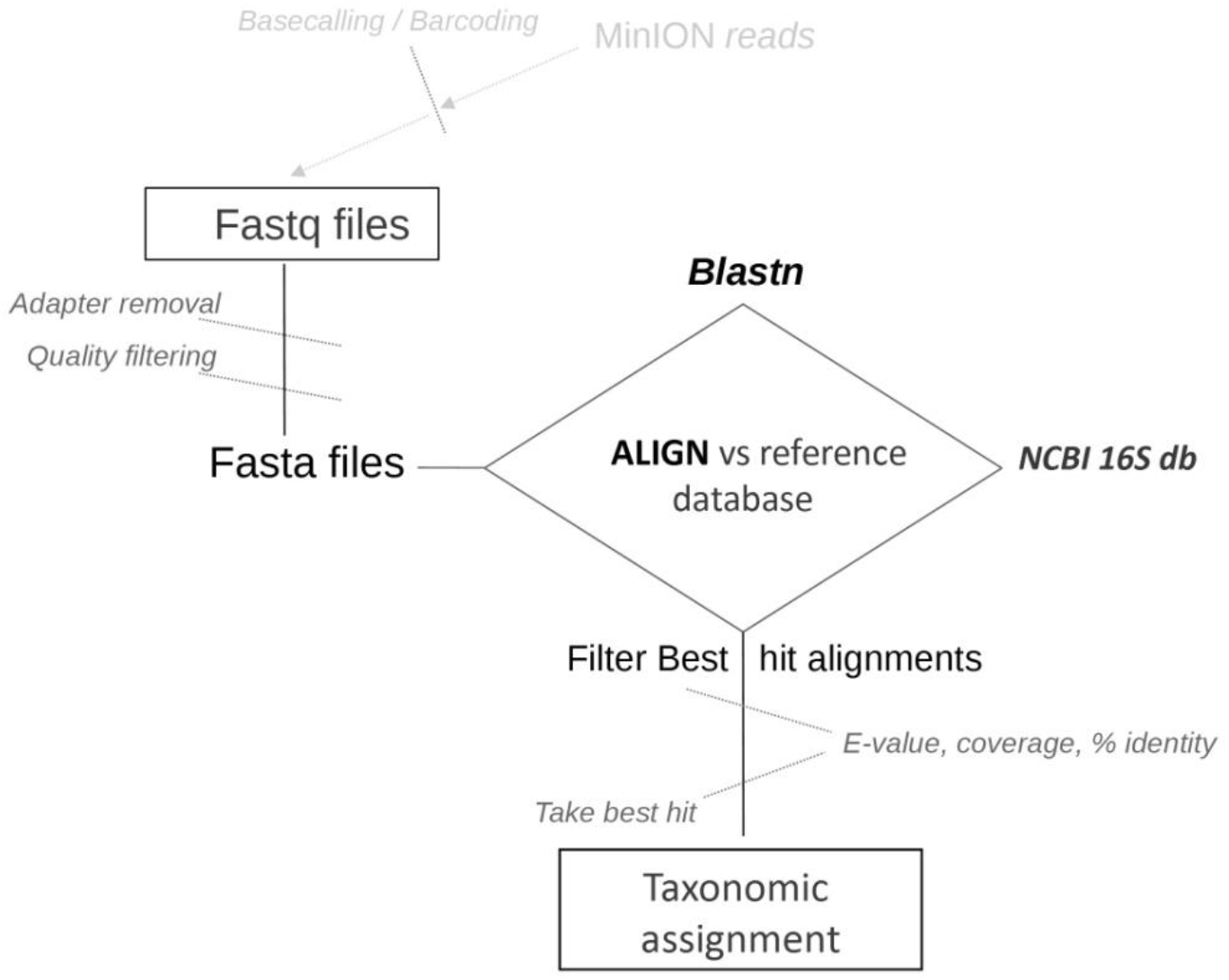
Scheme of the pipeline for taxonomic assignments.

**Figure 2.**
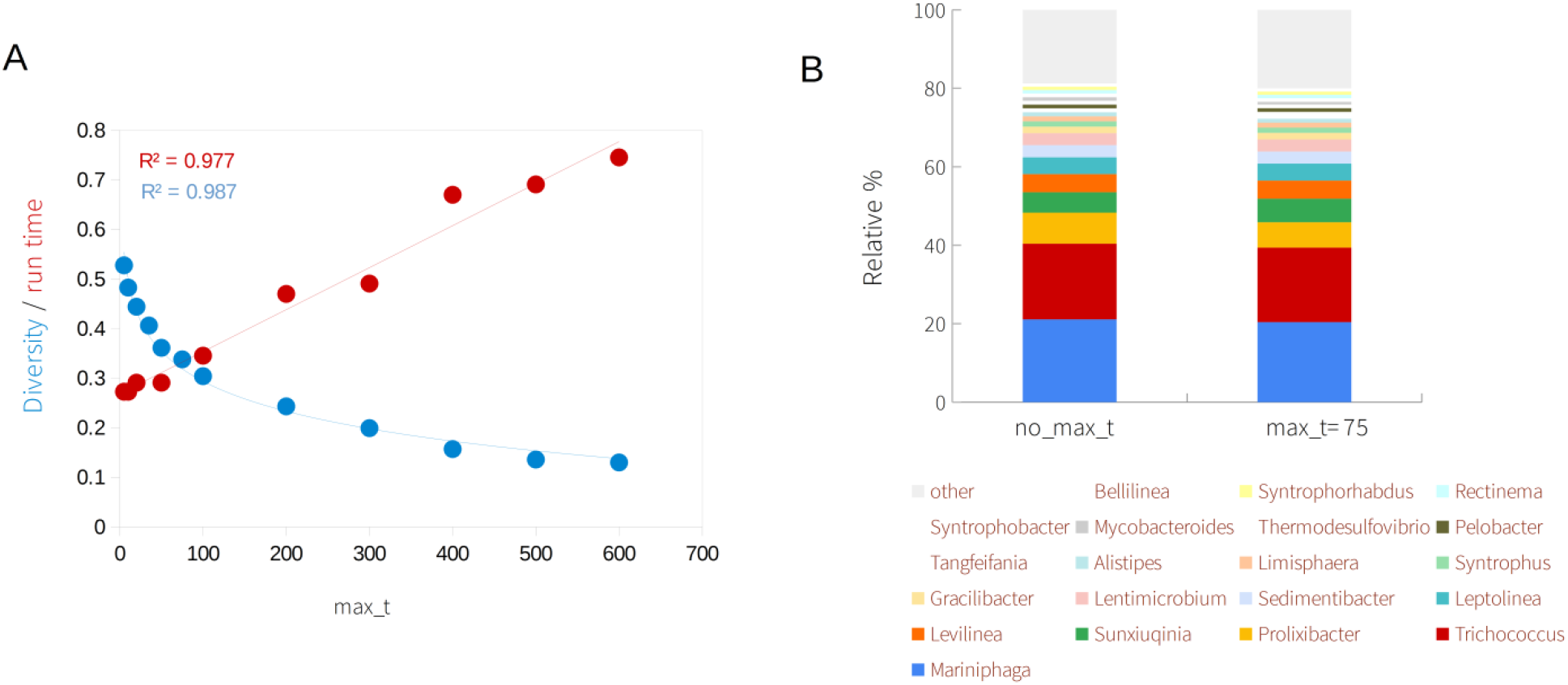
Effect of varying Max target (Blast option) on classification. Panel A: the effect of max_t on β-diversity (blue dots; non linear fit and correlation, with a function of the form f(x)=-log(kx)+w) and on run time (red dots; linear fit and R^2^ reported in red) is shown; max_t is set at different values, from 1 to 500 (the datapoint at max_t=600, refers to data for no_max_t option). Panel B: most abundant genera are shown, for no_max_t and for max_t=75.

The general structure of the pipeline (available at https://github.com/smm001/16S_ppm_public) is similar to ONT proprietary platform, Epi2me; however, aside from the differences in set up described specifically in the Results section, 16S_ppm allows for much more freedom (see Introduction, for the limitations of Epi2me) in customization, and the possibility to run it locally. The customization includes, together with all steps mentioned before (e.g. blast search options, such as max targets, e-value, coverage, etc; or quality filtering of reads), the possibility to choose if running the analysis on fastq inputs only (i.e. only performing classification; black bordered steps in Fig. 1), or fasta inputs, or running directly on sequencing data. Moreover, several configurable options can tailor the analysis: for once, the number of sequences to be classified can be set to specific values (default=20,000; more on this point in the Discussion) to allow, if needed, for rapid evaluations; overall, the analysis can be tailored to fit long and detailed calculations (e.g. on a server machine, also with high accuracy settings) or to fit agile and rapid evaluations (e.g. on a portable computer, for on-site performances). The latter being the main focus of this work, the default values in the configuration file are optimized for rapid scenarios. Scripting for pipelining, parsing, statistical evaluations were done with bash and Python (2.7 and 3.4). All calculations reported in this study were run on a portable workstation, with Linux Ubuntu (mainly, 14.04; tested also on 16.04), Processor Intel® Core™ i7-4900MQ CPU @ 2.80GHz × 8; RAM 32 GB; 2 × 256 GB SSD.

### Designed mutants

The strategy for designing mutants, with varying degrees of accuracy, was based on mimicking a distribution of accuracy level per each read, according to a Gaussian distribution centered on a specific value (the mean value for the distribution), and on a deviation (i.e. the width of the bell shaped curve). The probability density function is:

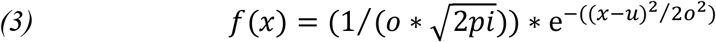

Where *u* is the mean, and *o* is the standard deviation. The estimated average level of accuracy for Nanopore reads with 16S gene is around 90% (*i*.*e*. slightly less than the average similarity for Nanopore reads in other applications, e.g. sequencing reads for human genomics). The most common variability was observed to range between 86% to 94%; for this study, it was chosen to simulate a profile of randomly mutated reads with *Eq. (3)*, with a variable *u* value and *o* = 2.8. This model was used to determine, for each reach a call for the accuracy (for example, if *u*=90 and *o*=2.8, the probability for each read was derived substituting these values in *Eq. (3)*).Once the accuracy is set for each read, the position of the mutated bases are randomly selected (random function, Numpy v1.4) within the complete fasta sequence, and the mutation type is also randomly selected (with each base being equally probable). Once all reads were mutated, the designed sample was written out in fasta format, labeled in accordance to its accuracy median value (*u*), for subsequent analyses (e.g. for Fig.3B; or for *S. rimosus*, in Table 2).

**Table 1.** Blast 2.2.31, with max target =1 and single (ST) or multi-thread (MT).

| Genus | max_t=1<br>MT, run1 | max_t=1<br>MT, run2 | max_t=1<br>MT, run3 | max_t=1,<br>ST, run1 | max_t=1,<br>ST, run2 | max_t=1,<br>ST, run3 |
| --- | --- | --- | --- | --- | --- | --- |
| Trichococcus | 162 | 152 | 158 | 156 | 156 | 156 |
| Mariniphaga | 86 | 90 | 90 | 86 | 86 | 86 |
| Prolixibacter | 34 | 34 | 34 | 34 | 34 | 34 |
| Leptolinea | 58 | 58 | 58 | 58 | 58 | 58 |
| Levilinea | 50 | 50 | 50 | 50 | 50 | 50 |
| Sunxiuqinia | 74 | 72 | 74 | 72 | 72 | 72 |
| Sedimentibacter | 26 | 26 | 26 | 26 | 26 | 26 |
| Tangfeifania | 44 | 44 | 44 | 44 | 44 | 44 |

**Table 2.** Analysis of *(i)* environmental sample, *(ii)* mock and *(iii)* mixtures of sample and mock.

| Genus | C3.1-20-8 | mB12 | C3.1-20-8<br>and mB12 | C3.1-20-8,<br>mB12 and<br>*S.rimosus* (A) | C3.1-20-8,<br>mB12 and<br>*S.rimosus* (B) |
| --- | --- | --- | --- | --- | --- |
| Mariniphaga | 4080 | 0 | 4080 | 4080 | 4079 |
| Trichococcus | 4053 | 0 | 4053 | 4053 | 4053 |
| Prolixibacter | 1292 | 0 | 1292 | 1292 | 1292 |
| Sunxiuqinia | 1255 | 0 | 1255 | 1255 | 1255 |
| Levilinea | 937 | 0 | 937 | 937 | 937 |
| Leptolinea | 909 | 0 | 909 | 909 | 909 |
| *Streptomyces* | 16 | 0 | 16 | 1013 | 1013 |
| Sedimentibacter | 703 | 0 | 703 | 703 | 703 |
| Lentimicrobium | 649 | 0 | 649 | 649 | 649 |
| **Halomonas** | 1 | 443 | 444 | 444 | 443 |
| Gracilibacter | 389 | 0 | 389 | 389 | 389 |
| Tangfeifania | 373 | 0 | 373 | 373 | 373 |
| Limisphaera | 297 | 0 | 297 | 297 | 297 |
| Syntrophus | 297 | 0 | 297 | 297 | 297 |
| Pelobacter | 207 | 0 | 207 | 207 | 207 |
| Alistipes | 205 | 0 | 205 | 205 | 204 |
| Syntrophobacter | 203 | 0 | 203 | 203 | 203 |
| **Marinobacter** | 35 | 165 | 200 | 200 | 200 |
| Rectinema | 199 | 0 | 199 | 199 | 199 |
| Syntrophorhabdus | 198 | 0 | 198 | 198 | 197 |
| Helicobacter | 197 | 0 | 197 | 197 | 197 |
| Thermodesulfovibrio | 196 | 0 | 196 | 196 | 197 |
| Geobacter | 193 | 0 | 193 | 193 | 193 |
| Flexilinea | 185 | 0 | 185 | 185 | 185 |
| Bellilinea | 185 | 0 | 185 | 185 | 185 |
| Smithella | 160 | 0 | 160 | 160 | 160 |
| **Psychrobacter** | 14 | 147 | 161 | 161 | 161 |

## RESULTS

### Blast versions and options

The overall structure of the pipeline is reported in Fig.1, and described in Methods. Some of the parameters of the pipeline requires particular attention; the first step was to assess the algorithm for similarity searches, Blast and its parameters. There are several options that can be set with Blast and that can affect the results. A particularly important one is the “max targets” option (max_t): if set, this option can limit the number of best hits considered by the algorithm, as it proceeds from the first alignments to the final hits; max_t can significantly reduce the burden of the calculations (time, but also size of the temporary files); however, it can also affect the results. Recently, a discussion emerged on this point (Shah et al, 2019; Madden et al, 2019): setting the max target parameter, particularly if set to its “quickest” (and thus time convenient) option, i.e. max_t=1, can return different results from the unrestricted search (as during intermediate steps, the algorithm may drop some legitimate candidates, which could otherwise be kept in consideration). This may be negligible in many cases, but it can affect results when many different, but similarly scoring, hits are present. Indeed, this may be the case with 16S genes; thus, considering the computational advantages it implies, the usage of the max_t option was specifically explored; additionally, at the light of the above mentioned discussion (Shah et al, 2019; Madden et al, 2019), two different Blast versions were considered: 2.2.31 (older, but still widely used) and 2.9.0, a recent version. Indeed, the tests with the older version showed interesting results: not only the max target option, but also the multi-thread option can affect the results (Table 1; data for a randomly selected subset of 1,000 reads from an environmental sample, SRP214990). Both these options would be important in the context of the application envisioned in this work, as they both make for significantly faster analyses. However, the repeatability of the analysis is a critical point: with Blast 2.2.31 (Table 1), if the multi-thread option (here, -num_threads=8) and the max_t=1 options are set, different runs return slightly different results. Interestingly, using max_t=1 with single thread results are repeatable (Table 1); similarly, if Multi-thread option is activated, but not the max_t option, results are consistent; thus, if both options are activated (as it is desirable, in this study) results are not repeatable with older Blast versions. In turn, Blast 2.9.0 did not present this issue: run with the same parameterization and to the same set of reads, it returned stable results (repeatable, in all cases). Therefore, the version 2.9.0 is recommended, at least for 16S related searched (and if employing the “max targets” flag with multi-threading).

To more closely examine the effect of max_t and multi-thread with Blast 2.9.0, a bigger scale analysis was performed on 10,000 randomly selected reads from an environmental sample with 16S ONT data (SRP214990). The analyses at different max_t values (all with multi-thread option activated, “-num_threads 8”) were evaluated as *(i)* diversity (inter-sample, with Bray Curtis diversity, BC) between results for different max_t values, and *(ii)* run time of the calculation, at different max_t values. Fig. 2, sums up the results; for a comparative visualization on the same graph, the data were normalized, from 0 to 1 (minimum/max values for diversity and run time are: 0/0.1 and 10 mins/60 mins, respectively): the variability rapidly decreases (non linear fit, blue line) with higher max target options, while run time increases linearly (red line). A good compromise is with max target set to 75 (max_t=75), for which results are both similar to the “unrestriced” Blast search (i.e. with no max target option, no_max_t) while significantly quicker. Additionally, it created significantly smaller temporary files (the size of intermediate Blast output are dependent on the number of targets considered, and can easily exceed few Gigabytes, for an input with thousands of sequences). In any case, the relative proportions of most abundant genera are well super-imposable (Fig. 2B; genera with relative abundance > 0.1%, for the “unrestriced” Blast, on the left; and for the max_t=75, on the right). Thus, the choice of max_t=75 seems to strike a good balance between time and performance: for example, with the reference equipment (see Methods) an analysis of 10,000 reads could be concluded in around 50 minutes, as opposed to 2 hours, for the unrestricted search with multi-thread (or escalate to over 13 hours for the unrestricted search, with single-thread analysis). Such a time gain could be important, for rapid on-site analyses. In the following paragraphs, all results refer to the employment of Blast v 2.9.0 with max_t=75 and multi-thread. As mentioned in the Methods section, the pipeline accompanying this work (16S_ppm) allows for ample customization, including running the search with smaller max_t values (e.g. max_t=5; for less precise, but very fast evaluations) or without the “max targets” option.

### Validation with experimental data

Typically, for preliminary test, a reference sample with a known composition is evaluated; these are referred to as mock samples. In this work, three types of validations with reference data are presented: *(i)* with a designed mock, for preliminary assessment, *(ii)* with two actual mocks (HM-783D, mock B12), and *(iii)* with a combination of mock and a real environmental sample data, to assess complex mixtures. The first test was done with the simulated mock. A mock made of 12 bacterial species (*Escherichia coli, Streptomyces rimosus, Streptomyces antibioticus, Bacillus subtilis*, P*seudomonas fluorescens, Rhizobium lusitanum, Flavobacterium johnsoniae, Flavobacterium capsulatum, Micromonospora echinospora*), each represented by 150 reads (for a total of 1800 reads; all covering the full length 16S gene) was created. Then, using an *in house* developed strategy for introducing random mutations (described in Methods), sets of designed mutant reads were created: briefly, for each of the 1800 reads of the mock, mutations were introduced to simulate a distribution of accuracy according to a specific value: for example, assuming a 90% accuracy, a random subset of mutations (no bases preferences; A, C, G, T all equally probable), scattered across the gene, was introduced, so that the average mutation rate per read was 10% (i.e. most reads will have an accuracy in the range 86 to 94% according to a distribution and parameterization described by *Eq (3)*; see Methods). With this approach, different mutant sets were created, for different degrees of accuracy: 98%, 95%, 92%, 90%, 85%, 80%. This test was meant to assess the sensibility of the pipeline to the degree of accuracy (here, for simplicity, assuming random mutations, with no significant bias within the 16S gene; this may not be exactly the case with ONT reads; however, low accuracy, rather than possible biases, is commonly referred to as the main limitation, for ONT data in 16S). Results are summed up in Fig. 3: panel A, reports the number of reads, actual and mapped, for each mock; panel B, shows the correlation between original and mutated data, as it varies with accuracy. The first observation is that accuracy impacts the results, with a non linear behavior. Grouping results by genus, Fig. 3B shows that for values above 90%, the mutation rate does not significantly affect the classification (R^2^≥0.97, for accuracy ≥ 90%); instead, for lower values results correlates less and less with the mock actual data (indeed, at 75% the Pearson R^2^ drop to circa 0.10). Therefore, with an expected accuracy similar to current Nanopore output, the Genus analysis seems reliable. At the species level results were less significant (as compared to Fig. 3B, R^2^< 0.90 in all cases, including the highest accuracy simulated). Nevertheless, if the actual 16S accuracy could increase to approach 95%, the upgrade would be certainly beneficial; at genus level, it should considerably reduce the number of artifacts (genus marked as “other” in Table of Fig. 3; these are the genera “created” by chance, by low accuracy introducing artificial richness in the sample); also, it should extend the analysis to a much more reliable species level (see also later paragraphs). The next step was to assess some “real” mock data. Two mocks were considered, the mock HM-783D (SRR8029996) and the mock B12 (SRR8351023): the first is a 16S metabarcoding dataset, while the second is a metagenomics dataset. For the mock B12: reads related to 16S gene were specifically extracted from the complete set of metagenomics reads, thus creating an *in house* derived dataset, composed of 16S reads (1,828 reads). Both mocks were then assessed with the 16S_ppm pipeline. The HM783D mock is composed of a mixture (at different concentrations) of 20 bacterial strains (of which 8 species, and 6 genera, with a relative abundance > 1%). Results are reported in Fig. 4 (left): all genera were found and the relative proportion of reads mapped correlated very well (R^2^ > 0.99) with the relative abundance in the sample (Cabral et al, 2017). At the species level the results were almost as good, with detection of all species (except *Propionibacterium acnes*), and a remarkable correlation between actual and calculated relative abundances (R^2^=0.961). The mB12 mock is composed of a mixture of 12 bacterial strains (*Thioclava sp. ES*.*032, Psychrobacter sp. LV10R520-6, Propionibacteriaceae bacterium ES*.*041, Muricauda sp. ES*.*050, Micromonospora echinofusca DSM 43913, Micromonospora echinaurantiaca DSM 43904, Micromonospora coxensis DSM 45161, Marinobacter sp. LV10R510-8, Marinobacter sp. LV10MA510-1, Halomonas sp. HL-93, Halomonas sp. HL-4, Cohaesibacter sp. ES*.*047*), belonging to 8 genera. Also in this case, the relative composition in the mock community is known, and different between different strains (e.g. see Fig. 2 of Sevim et al, 2019).

**Figure. 3.**
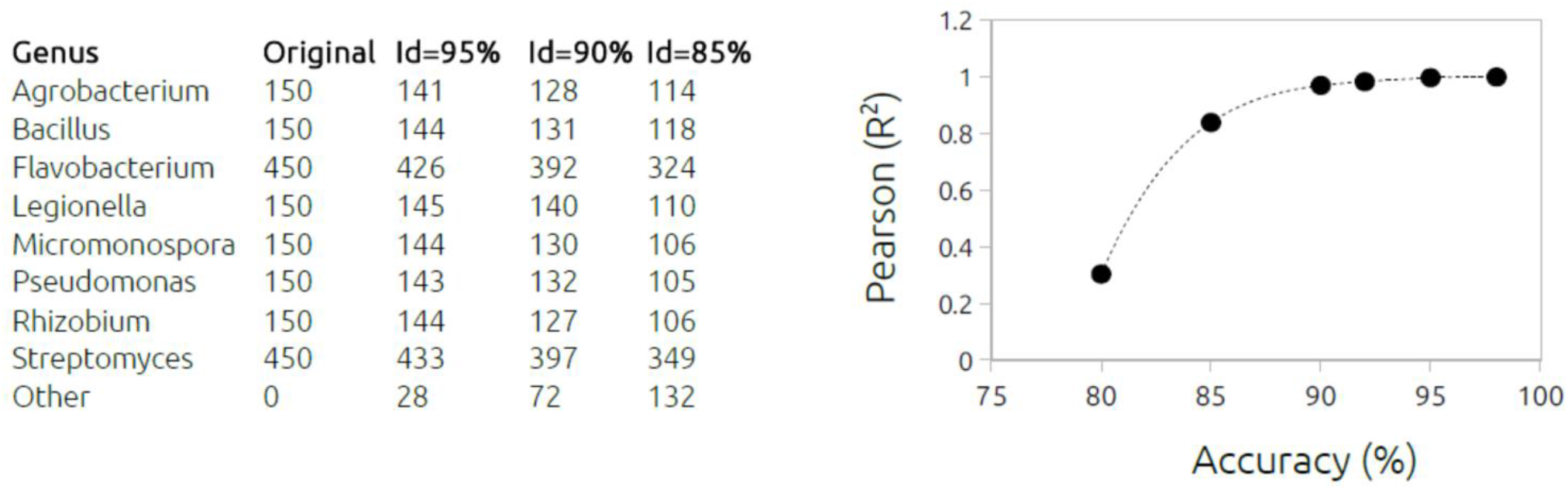
Analysis of a simulated mock: effect of average read accuracy on classification (genus level). On the left: reads matched to each genus, in different runs with varying %accuracy (Id=95%, 90%, 85%); on the right: plot of the correlation (Pearson,R^2^) between runs with varying % accuracy (from 80% to 97%) and the original (unmodified, Id=100%) data.

**Figure 4.**
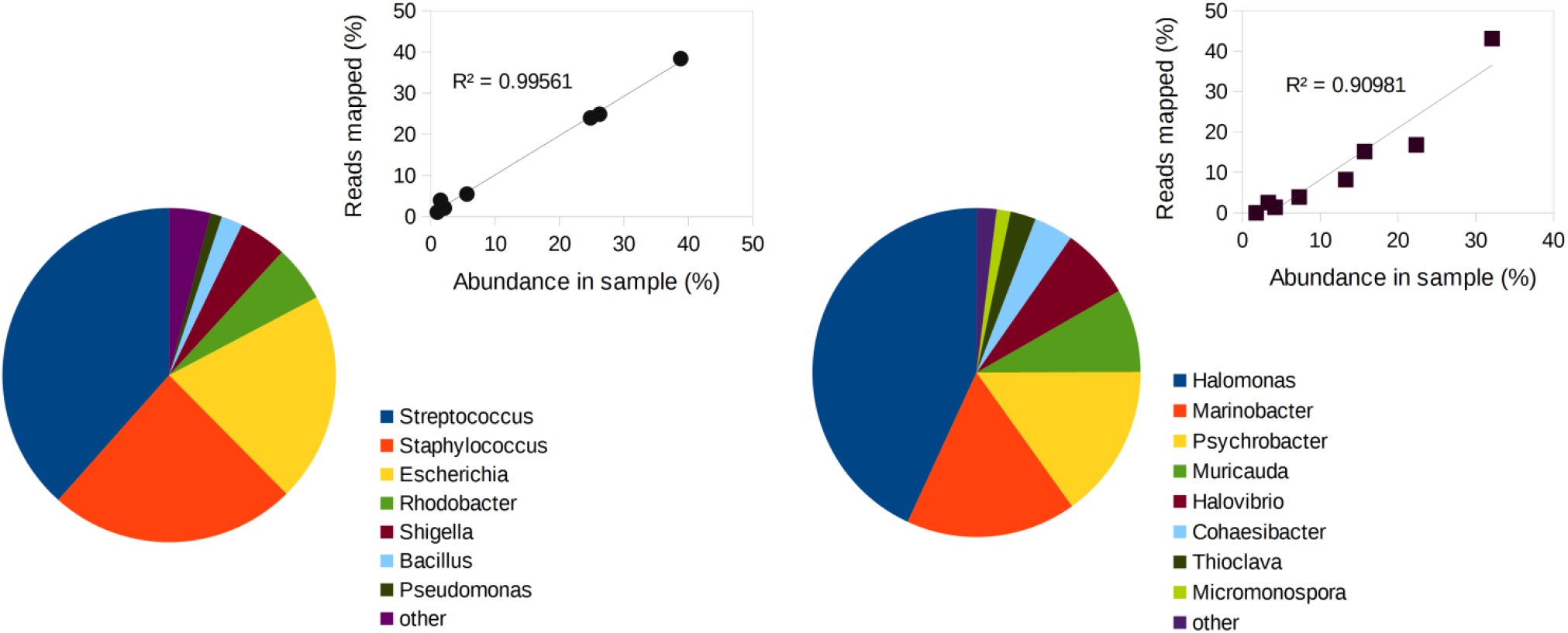
Analysis of bacterial mock B12). Pie chart of most abundant genera detected. In the upper-right section: correlation plot between the % reads mapped per each genus and the actual relative abundance (in mock); left: data for mock HM783D; right: data for mock B12.

After running the 16S_ppm scripts on this mock, all genera were found (Fig. 4, right), except for “Propionibacteriaceae bacterium ES.041”, which is not represented in the 16S NCBI db. Furthermore, the relative proportion of taxa detected by the pipeline correlated well with the relative abundances of the mock: plotting genus abundance (as relative abundance, expressed in molarity; Sevim et al, 2019) *versus* the reads mapped (as relative frequency) with 16S_ppm, provides a significant correlation (R^2^ =0.910; Fig. 4B). In this case, the analysis at the species level was not feasible, as several species in the mB12 mock (e.g. *Cohaesibacter sp. ES*.*047*; see the above list) are not present in the NCBI 16S reference database (and thus, could be not found). Altogether the analyses of both mocks provided satisfactory results, highlighting the capabilities of the pipeline to describe real data, at the genus level. To test the effect of a more complex mixture on the results, further analyses were done with a “real life” complex 16S sample: the same sequencing data (for an untreated anaerobic digester, sampled in Germany; project code SRP214990, run:SRR9695976, library:C3.1-20-8; 23,924 reads) previously considered (in Fig. 2A), was *(i)* analyzed “as it is” (“C3.1-20-8” in Table 2), *(ii)* spiked with 1,000 reads (randomly selected) from mock B12, (creating a mixed sample, with 24,924 reads, “C3.1-20-8 and mB12” in Table 2) and *(iii)* further spiked with 1,000 reads of *Streptomyces rimosus* (derived from the NCBI 16Sdb, and then mutated i*n silico*, with the approach described before), added to the previous mixture (creating a second mixed sample, “C3.1-20-8, mB12 and S.rimosus (A)” in Table 2). The 16S_ppm pipeline was run on these samples, and results are reported in Table 2 (genera with relative abundance > 0.5%): the outputs from the spiked samples (“C3.1-20-8 and mB12”; and “C3.1-20-8, mB12 and S.rimosus (A)”) accurately represent each specific change; in particular results at the genus level allow to trace all inputs from each modification, in the mixture. The experiment shows that the reads from different samples do not interfere with the classification: reads from mB12, once added to the set of reads for C3.1-20-8, do not “confuse” the classification of the latter sample. However, it should be noted that in a real life investigation of two samples, of which one is for the reference site (e.g. C3.1-20-8) and a second one represents a putative contamination (e.g. C3.1-20-8 with mB12 and *S*.*rimosus*), both samples are sequenced: thus, even assuming that both have identical material, minus the contaminants (mB12 and *S*.*rimosus*), the reads for C3.1-20-8 would be different for sample one and two, as the MinION sequences them independently. To simulate this effect, a variant of sample two was created with the following strategy. All reads in sample one (column “C3.1-20-8”, in Table 2) were substituted with the corresponding reference sequences of their best hits, taken from the NCBI 16S db (e.g. for read #1, its best hit with NCBI 16S db was selected, and its sequence assigned to read #1): this creates a new set of reads for C3.1-20-8 (more accurate, as they come from the reference database), while maintaining exactly the same taxonomic distribution. Then, these accurate reads were subjected to the same procedure for simulating lower accuracy reads described before (Figure 3, and Methods). This generates a new C3.1-20-8 sample, with similar accuracy (∼90%) to the first, and meant to simulate an independent run of sequencing for the same material (i.e. of the first C3.1-20-8 sample, second column in Table 2); then, the mocks were added to the mix, creating a second mixture sample, “C3.1-20-8, mB12 and *S*.*rimosus* (B)” (last column in Table 2). Results with this experiment confirmed, in all aspects, the “simpler” case (“C3.1-20-8, mB12 and *S*.*rimosus* (A)”, Table 2). Comparable results were obtained using a program for simulating Nanopore reads (Nanosim) (https://github.com/bcgsc/NanoSim;) to create ∼90% accuracy reads (data not shown). Of course, one has to further note that this is a simulated approximation, assuming an important simplification: even if sample two would be composed of the exact same material of sample one (minus the contaminants), a real life procedure would imply two separate 16S amplifications, for sample one and for sample two, before sequencing. This would create further variability; however this cannot be simulated accurately, and thus, in this example, this variability was ignored. In any case, the experiment described in Table 2 aims to approximate a real life situation, for a basic environmental analysis: a sample is taken in a control site (C3.1-20-8), and another (related) sample is taken for a putative contamination site (C3.1-20-8 with mocks). The application of ONT technology, with 16S metabarcoding and an appropriate bioinformatics pipeline, allows to trace the event of contamination within the polluted sample (e.g. see specific increase only in contaminating genera, in the last two columns of Table 2, as compared to the “uncontaminated” sample), clearly distinguishing the two sites. Similar analyses were conducted on additional samples (and mixture of mocks; see Supporting information), that supported results and conclusions drawn here. In the next section, the applicability of the approach to real world scenarios (with some additional data from environmental samples) is specifically discussed.

## DISCUSSION

A major advantage of ONT sequencing with MinION is portability, as well as the robustness of the protocols involved; the latter is not a minor feature: in recent years, several campaigns have been performed to test portable equipment and optimize reagents and protocols for molecular biology and sequencing library preparation, so that reliable operating procedures can be conducted in the field, including working in difficult conditions (Quick et al, 2016; Castro-Wallace et al, 2017; Arwyn et al, 2019). Indeed, with an affordable budget, for equipment and reagents, it is possible (and has been done, by more and more scientists and researchers) to perform a full investigation in the field, from sampling to data analysis (Maestri et al, 2019; Boykin et al, 2019). As for the latter, the approach outlined and validated in this study is specifically designed for rapid and affordable (here meant as low computational requirements, in terms of hardware and software, but also time and energy consumption) 16S metabarcoding, operated directly *in situ*, for environmental investigations, e.g. to check for contaminations or spills, including those putatively releasing pathogens in the environment. To put the work in perspective, with an estimate of the efforts involved, let’s describe a basic real life scenario, for an event similar to the one described in Table 2 (i.e. with a control site, and a putatively contaminated second site). The work flow would include: *(i)* sampling and DNA extraction (e.g. with one of the many commercial kits available), *(ii)* molecular biology protocol for 16S amplification (e.g. with ONT 16S metabarcoding kit), *(iii)* sequencing with MinION, connected to a (mid-range, see Methods) standard laptop, and *(iv)* bioinformatics (run on the same computer; with data analysis running, while the sequencing is ongoing) for sequencing and data analysis. The computational analysis can be scaled up at very different levels: for a “rapid evaluation” scenario 20,000 reads per sample could be sufficient, to obtain a quick but representative microbiological description of the system. In a similar “rapid scenario” case, the complete work-flow (with the pipeline parameterized as defined in the Results section), for two samples like those in Table 2, could take 7-8 hours of work (from DNA purification to data analysis; the lengthiest step being the PCR, for 16S metabarcoding), after which results (with taxonomy assignments) would be available.

In a typical case, this amount of reads/sample (20,000, for the “rapid scenario”) is sufficient for an informative description of the microbiome composition; if compared to more extended data (e.g. 200,000 reads), using 20,000 reads can be a very convenient approximation: it will accurately detect all the abundant taxa, and with good frequency approximation. To provide quantitative data in support to this claim, a representative case, based on real environmental data, is discussed. ONT 16S reads from the most recent (at the moment of this writing) experiment with MinION 16S barcoding (Gonçalves et al, 2020) were analyzed; this project (NCBI project: PRJNA598707) is well suited for assessing the effects of down-sampling, as it is characterized by sequencing runs with large outputs (in all cases > 200,000 reads for sample; to be noted, the project PRJNA554976, used for Table 2, did not lend well for this task, as it had a significantly lower output, < 50,000 reads/sample); in particular the data from the site called “Magallanes” (library:SRR10820646) were analyzed. The distribution of most abundant taxa (> 0.5%) are reported in Fig. 5. All genera are shared, and the correlation of the data generated for 20,000 reads with those generated for the extended dataset (250,000 reads) is very significant (R^2^=0.998). Moreover, the absolute values (i.e. of the relative abundance, for each genus) are very close, with an average deviation of 5%: given a relative abundance value as computed from the “rapid analysis” (“only” 20,000 reads), the expected variability is ±5%. The correlation remains significant for lower abundance genera (R^2^=0.921, for genera in the range 0.5% to 0.05%), albeit with a greater variability in absolute values (±16%). Indeed, 20,000 reads were sufficient to replicate the results highlighted in the original work (Gonçalves et al, 2020) for the most significant inter-samples differences. Further details and other data on this analysis, are reported in Supporting information (Fig. S3-S5). The effect of sub-sampling on diversity (at the genus level), was tested also with a randomly generated sample: starting from a random selection of 16S genes from the NCBI, 300,000 reads were simulated with Nanosim (https://github.com/bcgsc/NanoSim), thus creating a representative sample to be analyzed, for beta-diversity at different degrees of subsampling (from 1,000 randomly subsample reads, to 200,000 reads). For each level (1,000; 5,000; 10,000; 20,000; 50,000; 75,000; 100,000; 150,000, 200,000 reads), diversity was calculated against the reference (full) set of 300,000 reads, expecting diversity to decrease with increasing size of the subsample: results are reported in Fig. 5 (rightmost panel), and support the view that 20,000 reads (up to 50,000 reads) are a good practical choice for balancing rapid run times with results reliability (i.e. affinity with those for full scale calculations).

**Fig. 5.**
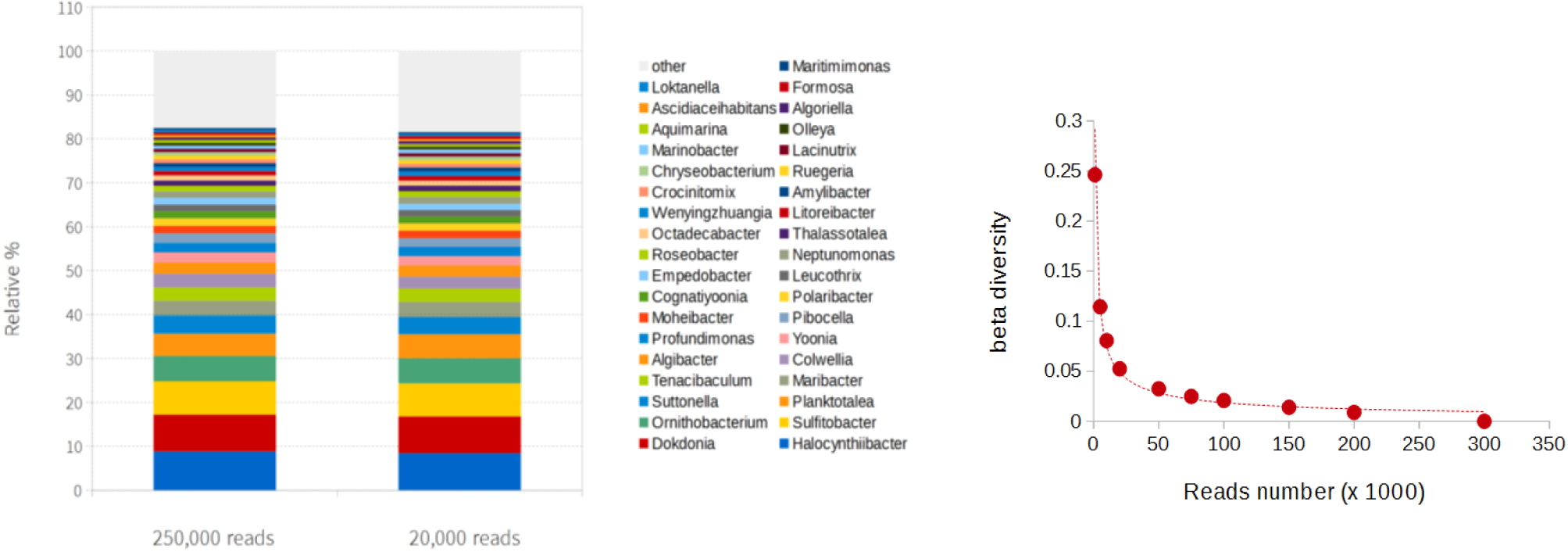
Effect of sub-sampling (at genus level). Panel on the left: genera in the “Magallanes” study with sub-sampled vs extended dataset; genera (> 0.1% abundance) are reported both for the complete analysis (left cumulative bar), and the sub-sampled (20,000 reads) analysis (right cumulative bar). Panel on the right: the effect of sub-sampling on beta diversity, in a representative sample.

Altogether, these results highlight that the “rapid evaluation” set up can be, at least for certain applications, a powerful tool. Considering the focus on rapid on-site work, the pipeline provided with this study runs with a default set-up for a “rapid evaluation scenario” (with search parameters as detailed in the Results section, and automatically limiting the analysis to 20,000 reads/sample). If a more detailed view is required (or, if the analysis is run without particular constrains, e.g. on a server with no specific limitations), the pipeline configuration can be tailored accordingly. Indeed, a deeper coverage (than 20,000 reads per sample) is required in various cases, particularly if some very low abundance taxa are specifically addressed (e.g. to check for their presence, albeit in small amounts); in any case, extending it to > 150,000-200,000 reads per sample, may not necessarily provide a good cost/benefits balance (as reported in the original work; see also Supporting information, Fig. S3, S4).

In conclusion: for a on-site analysis, a good compromise could be to first analyze the data at 20,000 reads per sample (“Rapid analysis”), while the MinION sequencing is still running: for two samples, aiming for 20,000 (fastq) reads/sample, an indicative time frame could be to start the analysis 30-40 minutes after the sequencing have started (or as soon as a sufficient number of fast5 reads is available). This rapid analysis will provide a quick, but already indicative response (as the one in Table 2; or in Fig. 5; and in Supporting information, Fig. S3, S5); meanwhile the sequencing would continue, for example targeting 100,000 to 200,000 fast5 reads per sample. A subsequent analysis, e.g. at 150,000 reads/sample, will take many hours in excess of the rapid analysis (altogether, making it more suitable for an overnight calculation). Naturally, in cases of multiple (i.e. > 2) sampling sites, the time taken for the complete analysis increases accordingly. Indicatively, for 6 barcodes, a complete “Rapid analysis” could take a full day of work (from sampling to results), while it could extend to some days if more reads (≥ 100,000/sample) are analyzed.

## Data availability

The scripts for the pipeline are available at https://github.com/smm001/16S_ppm_public

