## Supporting_information for "A computational strategy for rapid on-site 16S metabarcoding with Oxford Nanopore sequencing"

Additional material for the paper “A computational strategy for rapid on-site 16S metabarcoding with Oxford Nanopore sequencing”, Stefano M. Marino,

### SUPPORTING INFORMATION

*Analysis with ZymoBIOMICS™* - An additional validation was done with a third reference sample, the mock ZymoBIOMICS™ (Zymo). The mock is composed of *Pseudomonas aeruginosa*, *Escherichia coli*, *Salmonella enterica*, *Lactobacillus fermentum*, *Enterococcus faecalis*, *Staphylococcus aureus*, *Listeria monocytogenes*, *Bacillus subtilis*, as well as two yeasts (*Saccharomyces cerevisiae*, *Cryptococcus neoformans*). The reads were downloaded from the Loman lab (https://lomanlab.github.io/mockcommunity/ library Zymo-GridION-EVEN-BB-SN-PCR-R10HC-flipflop); this is a metagenomics experiment: first the dataset was “filtered” for the extraction of only those reads containing 16S gene sequences (partial or full), similarly to what done for the mock B12, described in the main text. This created a dedicated Zymo 16S dataset of 7096 reads, which was then fed to the 16S_ppm pipeline, for analysis. The relative abundance of the species in the mock is: *Pseudomonas aeruginosa* (4.2), *Escherichia coli* (10.1), *Salmonella enterica* (10.4), *Lactobacillus fermentum* (18.4), *Enterococcus faecalis* (9.9), *Staphylococcus aureus* (15.5), *Listeria monocytogenes* (14.1), *Bacillus subtilis* (17.4) (Zymo Research Corp Instruction Manual ZymoBIOMICS; Gonçalves et al, 2020). All genera in the mock were detected (Fig. S1), with a sufficient correlation between computed and estimated relative abundances ($R^2$=0.59). This value is very similar and confirms what previously reported for the same sample, with the ONT platform Epi2me (Gonçalves et al, 2020, including data from Supporting information). At the species level the detection is the results are less satisfactory (correlation, $R^2$=0.44): all species are found, but with some misclassifications (e.g. *Escherichia fergusonii* in place of *Escherichia coli*, to which only less than 1% of overall reads are assigned; and *Listeria innocua* or *Listeria welshimeri*, as the most abundant species of the Listeria genus, instead of *Listeria monocytogenes*).

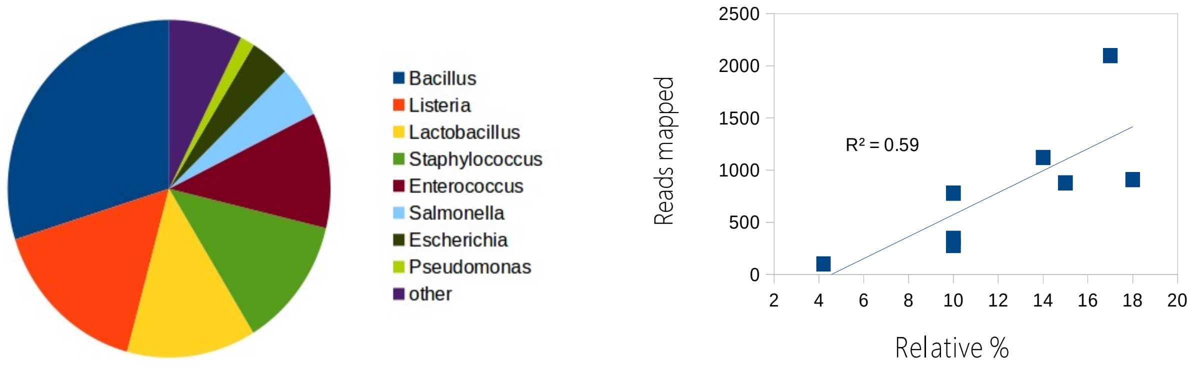


**Figure S1. Analysis of the Zymo mock.** On the left, pie chart with relative abundances of genera detected. On the right side of the picture, the correlation (with Pearson $R^2$ reported) between the number of reads mapped and the actual relative abundance of the *ZymoBIOMICS™ mock.*

The Zymo mock was then used to create analyses with *ad hoc* variations (i.e. designed mocks, to assess changes; similarly to what done in the main text for mock B12). The Zymo mock was modified in 3 samples: *(i)* with *S.rimosus* reads (mutated with 90% accuracy; see procedure described in the main text), *(ii)* with all reads for Halomonas genus (788 reads) taken from the mock B12, and *(iii)* with 1000 randomly selected reads, also from the mock B12. Results are summed up in Table S1, which shows that the modifications (the spiked material) do not interfere with the original sample. As in the similar case analyzed reported in the main text, also in this test it was possible to clearly trace the contribution coming from the different "external additions", for each of the mixed samples (see last column), with no significant effect of cross-sample "confusion" of the data (which, thus, remains traceable in each specific contribution to the mock).

**Table S1. Top genera for Zymo mock** as it is and with different spikes

| GENUS | Z, Zymo * | Z + *S.rimosus* | Z +Halomonas (mB12) | Z + mB12 (1000 reads) |
|---|---|---|---|---|
| Bacillus | 2098 | 2098 | 2098 | 2098 |
| Listeria | 1121 | 1121 | 1121 | 1121 |
| Lactobacillus | 908 | 908 | 908 | 908 |
| Streptomyces | 0 | **907** | 0 | 0 |
| Staphylococcus | 879 | 879 | 879 | 879 |
| Enterococcus | 778 | 778 | 778 | 778 |
| Halomonas | 1 | 1 | **789** | **444** |
| Salmonella | 346 | 346 | 346 | 346 |
| Escherichia | 278 | 278 | 278 | 278 |
| Marinobacter | 0 | 0 | 0 | **164** |
| Psychrobacter | 1 | 1 | 1 | **148** |
| Pseudomonas | 102 | 102 | 102 | 103 |

Taken together, results support the applicability of the approach to complex mixtures, e.g. for tracing contaminants in different samples. Finally, the analysis was done with a real life complex sample (as for Table 2, main text), spiked in with the reads from the Zymo mock (Table S2). Also in this case, results shows no interference, hence the approach was capable of working with spikes, to evaluate the

**Table S2. Most abundant generea, for an actual environmental sample**, with or without spikes (see text).

| Genus | C3,1 + Zymo | C3,1 | Zymo |
|---|---|---|---|
| Mariniphaga | 4080 | 4080 | 0 |
| Trichococcus | 4054 | 4053 | 0 |
| Bacillus | 2169 | 71 | 2098 |
| Prolixibacter | 1292 | 1292 | 0 |
| Sunxiuqinia | 1255 | 1255 | 0 |
| Listeria | 1122 | 1 | 1121 |
| Levilinea | 937 | 937 | 0 |
| Lactobacillus | 935 | 27 | 908 |
| Leptolinea | 909 | 909 | 0 |
| Staphylococcus | 882 | 3 | 879 |
| Enterococcus | 794 | 16 | 778 |
| Sedimentibacte | 703 | 703 | 0 |
| Lentimicrobium | 649 | 649 | 0 |
| Gracilibacter | 389 | 389 | 0 |
| Tangfeifania | 373 | 373 | 0 |
| Salmonella | 346 | 0 | 346 |
| Syntrophus | 297 | 297 | 0 |
| Limisphaera | 297 | 297 | 0 |
| Escherichia | 278 | 0 | 278 |
| Pelobacter | 207 | 207 | 0 |
| Alistipes | 206 | 205 | 0 |
| Syntrophobacte | 203 | 203 | 0 |

effects of a perturbation on a complex, real-life sample (the test was meant to mimic a basic situation of: control site *versus* control with contamination, e.g. from a spill, to test if the pipeline can trace the contamination in a mixed sample).

*Mock analysis with LAST aligner* – Comparisons were made between different aligners, Blast and LAST, using the three mocks described in this work. LAST was used both in default configuration, and with customized parameters (as previously employed with ONT data, Xia et al, 2017).

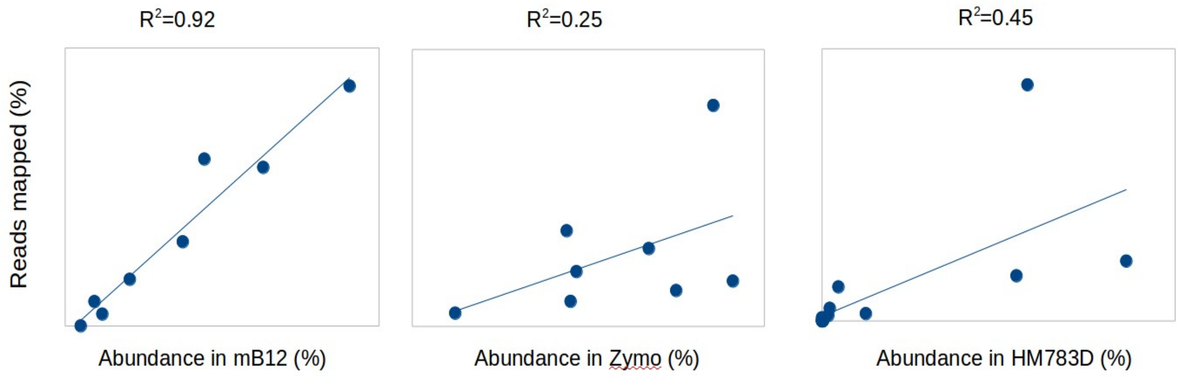


**Figure S2. Correlation of data with a different aligner with mocks.** Correlation of the % of mapped reads to each genus in a mock plotted against the actual abundance of that genus in the sample. Data refers to the LAST aligner (version 9-79, with default parametrization).

In 2 out of 3 cases (Fig. S2, and compare with Fig. S1, and main text Fig. 4), LAST results proved less effective than those for the corresponding Blast analysis; instead, for mock B12, its performances was on par (slightly better) with that of Blast. However, in all cases LAST returned a significantly lower amount of annotated reads (~30-35% of reads were unclassified, compared to ~1-2% of Blast based analysis). LAST was also tested with custom parametrization (see reference Xia et al, 2017, in main text); results did not improve, while the custom scheme of parameters slowed down the search, and caused a much heavier computational load: for example, with mock B12, the correlation was slightly lower (to the one with default, Fig. S6), with a Pearson $R^2$=0.87, but at the cost of a ~10 times higher run time (and with a similar increase in space, with temporary files, e.g. maf files, in the range of a few Gigabytes). While LAST, especially with default parameter, can still be considered a viable option, in our tests and for our tasks its performances were less effective than those of Blast.

*Effects of subsampling on 16S metabarcoding with Nanopore* – While preparing this manuscript a study (project id: PRJNA598707) was published reporting a 16S rRNA gene metabarcoding analysis of microbiota variability in parasites affecting salmons, sampled in different locations in South America (Gonçalves et al, 2020**)**. The study provided a large output (> 200,000 reads, for all samples) and, thus, it was employed for studying the effect of sub-sampling on the taxonomic description of a sample. In particular, the data for the site called “Magallanes” (library: SRR10820646; 250,000 reads were employed). Sub-sampling was done at 1,000, 5,000, 10,000, 20,000, 50,0000, 75,000, 100,000 and 150,000 reads; then, each sub-sampled set was given in input to the pipeline. Results were compared at genus level, for diversity: *(i)* intra-taxon (α-diversity, with Shannon) and *(ii)* inter-taxa (each subsample compared with the complete set of reads, evaluated with β-diversity, Bray-Curtis, BC, metrics). Furthermore, the sub-sampling was used to to assess the effect of the number of reads on computational time. Fig. S3 shows the α-diversity (blue dots; normalized for the max value, 4.05) and the β-diversity (red dots, reporting actual BC values; each value reflects the comparison between *i-th* subsampling and the complete dataset). In both cases, the values approached those of the complete dataset (> 200,000 reads) at around 20,000 reads.

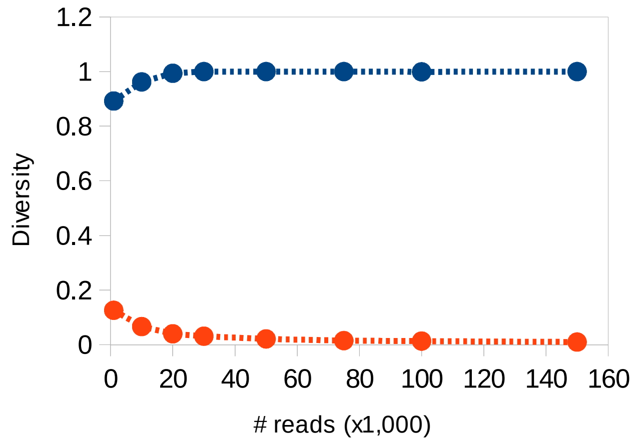


**Figure S3. Effects of subsampling on α- and β- diversity.** The "Magallanes" (SRR10820646) study, described in the main text (Fig. 5) is analyzed. This library produced > 600,000 reads (after quality filtering, see Methods). Here, 250,000 reads were used (for bigger values, diversity does not increase, while the computational burden increases considerably). The α-diversity is reported in blue (to be plotted in the same graph, values are normalized to its max), β- diversity in red.

The running time, as it varies with different subsets is reported in Fig. S4; the difference between analyzing 20,000 or 250,000 reads is very significant; it may not be advisable to run the full dataset, if the overall composition and diversity in a sample is sought after (Fig. S3 vs Fig. S4), e.g. for rapid detection of a contaminant or a significant spill between samples.
Naturally, as discussed in the main text, for a more detailed description (particularly, to assess taxa with very low abundances, if that is the specific target of the investigation), one can run (alongside the rapid test, with 20,000 reads) a more exhaustive (e.g. > 100,000 reads) analysis.

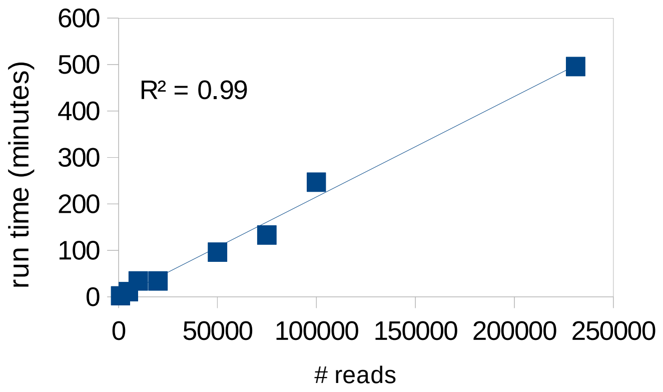


**Figure S4. Effects of subsampling on comutational run time.**

The PRJNA598707 project, including the above SRR10820646 and other six samples (for other sites: SRR10820647, SRR10820648, SRR10820649, SRR10820650, SRR10820651, SRR10820652), was also employed to assess the differences, highlighted in the accompanying publication (Gonçalves et al, 2020), between samples, at the genus level; the goal was to compare some of the results of the original work (derived from analysis of the full dataset of reads) with those derived here by the analysis of only 20,000 reads per sample (to assess and evaluate its feasibility, for a rapid response). In the work of *Gonçalves et al*, four genera were particularly highlighted, for the differences showed across different sites and samples, as well as for their biological relevance.

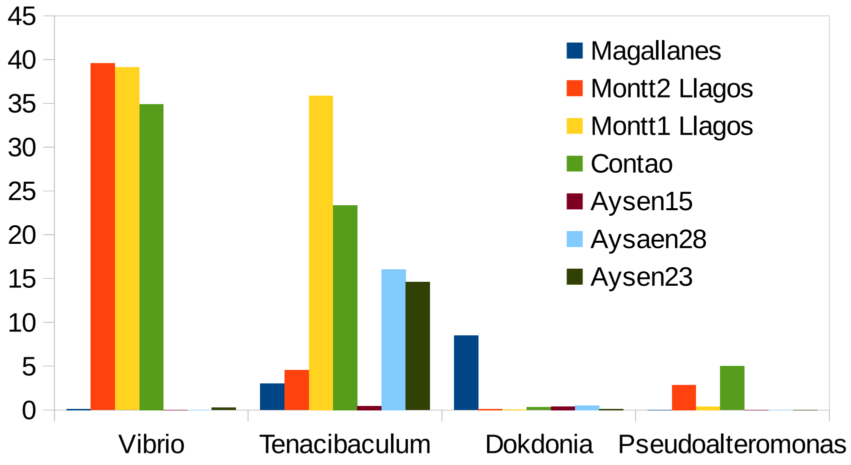


**Figure S5. Relative abundance of some relevant genera, highlighted in PRJNA598707 study.**

These genera, and their distribution (as relative frequency, i.e. relative % of reads in the sample) in all seven sample are reported in Fig. S5, based on the “rapid evaluation scenario” (20,000 reads/sample, with the 16S_ppm pipeline, set up as described in the Results section). The results replicated well (both as relative proportions and magnitude) the data reported in the original study (compare them with Fig. 5C, of Gonçalves et al, 2020). Furthermore, considering other differences highlighted in the original publication, the data with 20,000 reads are sufficient to detect main trends in the the data: for example, the most important (for size and significance) difference between the Magallanes site (Fig. 5B, of Gonçalves et al, 2020), and the average for sites in Los Lagos, are correctly detected (Dokdonia, 8.5% in Magallanes, 0.2±0.12 % in Los Lagos; Octadecabacter , 1.1% in Magallanes, 0.02±0.02 in Los Lagos).
